# Developmental links between play behavior and brain network integration

**DOI:** 10.64898/2026.03.26.714609

**Authors:** Monami Nishio, Maayan Ziv, Monica E. Ellwood-Lowe, Juan Ignacio Sanguinetti, Solange Denervaud, Kathy Hirsh-Pasek, Roberta M. Golinkoff, Allyson P. Mackey

**Affiliations:** Department of Psychology, School of Arts and Sciences, University of Pennsylvania, Philadelphia, US; Graduate School of Education, Stanford University; CIBM MRI imaging and technology, Polytechnical School of Lausanne, Swiss Federal Institute of Technology Lausanne (EPFL), Lausanne, Switzerland; Department of Psychology, Temple University, Philadelphia, US; Brookings Institution, Washington, US; School of Education, University of Delaware, Newark, US

**Keywords:** play, creativity, default mode network, salience network, control network, functional connectivity, Montessori

## Abstract

Play is a fundamental aspect of childhood and plays a crucial role in the development of creativity, yet its neural mechanisms remain poorly understood. We tested the hypothesis that more frequent play is associated with stronger functional integration among the default mode network (DMN), executive control network (CN), and salience network (SAL), as these cortical networks have been implicated in creativity in adults. In a preregistered study of infants and toddlers (Study 1; *N* = 143, 10 months–3 years, 67 boys, Baby Connectome Project), parent-reported play and imitation behaviors increased sharply from 1 to 2 years, and were associated with stronger within-DMN connectivity and DMN-CN coupling, controlling for age, sex, and head motion. In middle childhood (Study 2; *N* = 108, ages 4–11 years, 52 boys), parent-reported play frequency declined with age, as did cross-network coupling involving SAL. However, children who engaged more frequently in play showed higher DMN-SAL and CN-SAL connectivity. Finally, in a quasi-experimental comparison (Study 3; *N* = 45; ages 4-12 years, 20 boys), children enrolled in a curriculum that includes guided play (Montessori) showed higher DMN-SAL and DMN-CN connectivity than peers in traditional schools, suggesting that pedagogies that center child-led exploration might enable protracted brain network integration. Across these three studies, play was consistently associated with greater integration among DMN, SAL, and CN, a pattern previously linked to creativity in adults. Our findings offer a potential mechanism linking childhood play to later creativity through its role in supporting brain integration during development.

**Public Significant Statement:**

- Play is widely believed to nurture children’s creativity, yet the brain mechanisms behind this link are not well understood.
- Across three studies from infancy to middle childhood, we found that more frequent play was associated with stronger integration among brain networks tied to imagination, attention, and control.
- These findings suggest that play may help build the neural foundation for later creative thinking.

## Introduction

Play is widely considered to be critical for children’s cognitive development (Gilpin et al., 2015; Jaggy et al., 2023; Mireille Smits-van der Nat et al., 2024; Muentener et al., 2018; Stagnitti & Lewis, 2015; White et al., 2021). Children who engage in more play tend to have stronger executive functioning skills (Muentener et al., 2018; White et al., 2021), socioemotional skills (Gilpin et al., 2015; Jaggy et al., 2023; Mireille Smits-van der Nat et al., 2024), and language development (Stagnitti & Lewis, 2015). One of the most consistently documented skills associated with play is creativity (Burgdorf & Panksepp, 2006; Golinkoff et al., 2006; Hoffmann & Russ, 2012; Luo et al., 2024; Moore & Russ, 2008; Mullineaux & Dilalla, 2009; Russ 2014; Russ, 2020; Russ et al., 1999; Russ & Wallace, 2012; Siviy, 2016; Wallace & Russ, 2015; Yogman et al., 2018), the ability to generate novel and useful ideas (Runco & Jaeger, 2012; Skene et al., 2022). Play is not only correlated with creativity, it also been shown to causally enhance creativity (Howard-Jones et al., 2002; Moreau & Engeset, 2016): Children given the opportunity for free play created more imaginative and colorful drawings than children who received direct instruction (Howard-Jones et al., 2002). Why does play enhance creativity? One explanation could be that play creates a more open mindset, changing neural dynamics (Siviy, 2016; Weisberg et al., 2014). Here, we test the hypothesis that repeated engagement in play is associated with the development of neural networks implicated in creativity.

Across multiple studies, three large-scale brain networks, the default mode network (DMN), executive control network (CN), and salience network (SAL) have been consistently implicated in creativity in adults (Beaty et al., 2014, 2016, 2018; Hoffmann & Russ, 2012; S. Lee et al., 2021). The DMN is involved in processing everything we cannot sense directly, including the past, the future, and the minds of other people (Buckner et al., 2008; Buckner & Carroll, 2007). It is involved in imagination: generating ideas that are not, or are not yet, true (Buckner & Carroll, 2007). The CN plays a key role in setting goals and selecting actions (Miller & Cohen, 2001). The SAL is concerned with the here and now: spotlighting what is important and filtering out distractions (Seeley et al., 2007). The interactions among these networks can be quantified with resting-state fMRI: regions that frequently work together show positive correlations in fMRI signal, while regions that suppress each other are negatively correlated. In adults, the DMN is typically negatively correlated with both the CN and SAL (Fox et al., 2005), consistent with the idea that there is a tradeoff between attention to the internal world and attention to the external world (Buckner et al., 2008; Fox et al., 2005). These anticorrelation get significantly stronger over the course of childhood (Tooley et al., 2022). However, highly creative adults show a correlation structure more similar to that of children: DMN is less segregated from both CN and SAL (Beaty et al., 2014, 2016, 2018; Hoffmann & Russ, 2012; S. Lee et al., 2021). If play boosts creativity, then it may do so by maintaining connections among these networks.

Play encompasses a wide variety of behaviors, and the frequency of these behaviors varies over the course of childhood (Lillard et al., 2013; Sidhu et al., 2022; Yogman et al., 2018). Sensorimotor play involves touching and manipulating objects. In infancy, this play may be primarily about discovering the sensory properties of objects and their motor affordances, but later in childhood, this could involve more goal-directed forms of play, such as stacking blocks to build a tower (Lillard, 2015). Sensorimotor play is important for the development and coordination of sensory and motor skills (Lillard, 2015). It also provides a primary context for exploratory learning about object properties and action-outcome contingencies (Sidhu et al., 2022). Symbolic pretend play involves simple object substitution, such as using a block as a car (Bergen & Mauer, 2000), while dramatic pretend play includes more complex social role-playing, such as assigning characters and acting out roles (Hoffmann & Russ, 2012; Russ, 2020), which requires taking on the perspective of another. Play can be either independent or social and is often guided by adults to support learning while still preserving children’s agency (Skene et al., 2022; Weisberg et al., 2016).

We can combine what we know about the developmental trajectories of cortical networks with changes in play behavior to create a hypothetical framework for their interplay (Figure 1). From birth to around 1.5 years, infants engage primarily in exploratory sensorimotor play (Herzberg et al., 2022). At this age, sensorimotor networks change rapidly, as these regions develop relatively early (Gao, 2025). From around 18 months to three years of age, children transition to pretend play (Campbell et al., 2018), a shift that likely depends on the maturation of the DMN. During childhood, unstructured free play tends to decline, although some forms of play or playful learning may follow different developmental trajectories (Hirsh-Pasek et al., 2010; Hofferth & Sandberg, 2001). In parallel with declines of free play, connectivity among cortical networks differentiates: DMN–CN coupling shifts from positive to the characteristic adult anticorrelation (Yin et al., 2025), and competitive interactions between DMN and SAL emerge (Yin et al., 2025).

**Figure 1.**
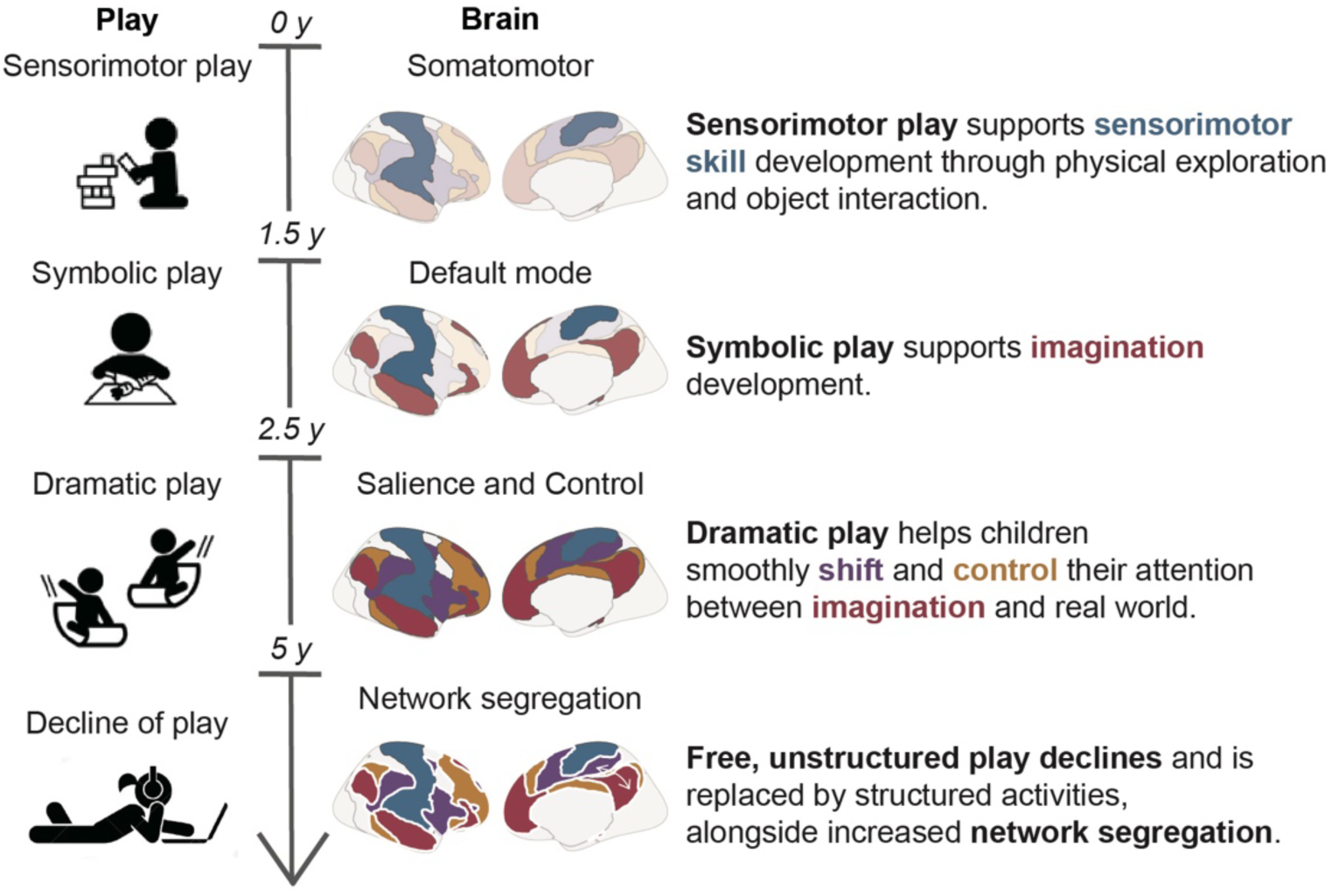
Hypothetical framework linking play and brain functional network development.

Parallels between the development of play and changes in network dynamics raise the possibility that play not only reflects, but may also contribute to the maturation of functional networks. The integrative nature of pretend play, requiring flexible transitions between imagination and external cues (Hoffmann & Russ, 2012; Russ, 2020; Russ & Wallace, 2012), could foster dynamic coupling among DMN, CN, and SAL. Notably, although greater integration among these networks has been linked to creativity, certain cognitive functions depend on network segregation (Kelly et al., 2008; Fox et al., 2005; Owens et al., 2020). For example, the anticorrelation between the DMN and SAL is thought to be essential for efficient goal-directed performance (Kelly et al., 2008). Thus, the flexible modulation between segregated and integrated network states may be particularly adaptive (Beaty et al., 2018).

Here, we examined associations between play experience and functional network development. In Study 1, we tested the hypothesis that higher levels of play behavior in infancy would be associated with stronger within-network connectivity among the DMN, and greater functional coupling between the DMN and the CN, beyond age-related changes in network organization. In Study 2, we tested whether age-related changes in connectivity within and between the DMN, CN, and SAL corresponded to declines in play behaviors observed during childhood. In Study 3, we examined whether children attending Montessori schools, educational contexts that emphasize self-directed, exploratory, and hands-on learning (Marshall, 2017; Rand & Montessori, 1941), showed greater functional network integration than peers in traditional schools. Across all studies, we controlled for socioeconomic status (SES), given its known associations with the nature and frequency of children’s play (Clearfield et al., 2014; S. Li et al., 2022; Mohan & Bhat, 2022; Rubin et al., 1976). Together, these studies test the hypothesis that play is linked to the integration of cortical networks throughout development.

## Study 1

In Study 1, we examined how play behavior in infancy relates to patterns of connectivity among major brain networks.

### Method

This study’s design and hypotheses were preregistered; see https://osf.io/42qru/overview?view_only=483cccb179f94ac98b4295df9d1fcfbe. ***Participants***

The Baby Connectome Project (BCP) dataset (https://nda.nih.gov/edit_collection.html?id=2848) includes 343 infants and 812 scans (Howell et al., 2019). Participants were screened to exclude infants born before 37 weeks of gestation, weighing less than 2000 g, or experiencing significant delivery complications (e.g., neonatal hypoxia, NICU stay > 2 days). Additional exclusions applied to children who were adopted, had a first-degree relative with autism, intellectual disability, schizophrenia, or bipolar disorder, presented with significant medical/genetic conditions affecting development, or had contraindications for MRI. Children were also excluded if their mothers had pre-eclampsia, placental abruption, HIV, alcohol/drug use, or were unable to consent in English. After quality control procedures on structural and functional data, the final BCP sample included 458 resting-state fMRI scans from 222 infants. Of these, 143 infants had measures of play behavior. The participants’ demographic characteristics are presented in Table 1.

**Table 1.** Demographics of participants.

|  | **Study 1**<br>**(N = 143)** | **Study 2**<br>**(N = 108)** | **Study 3**<br>**Montessori**<br>**(N = 22)** | **Study 3**<br>**Traditional**<br>**(N = 23)** |
| --- | --- | --- | --- | --- |
| **Age (years)** | 1.910 ± 0.6090 | 6.729 ± 1.382 | 8.567 ± 1.869 | 8.564 ± 1.826 |
| **Sex (male)** | 67 (47%) | 52 (48%) | 11 (50%) | 9 (39%) |
| **Ethnicity** |  |  |  |  |
| Hispanic or Latino | 11 (8%) | 17 (16%) | 0 (0%) | 0 (0%) |
| No Response | 0 (0%) | 1 (0%) | 22 (100%) | 23 (100%) |
| **Race** |  |  |  |  |
| White | 117 (82%) | 45 (42%) | 22 (100%) | 23 (100%) |
| Black | 2 (1%) | 61 (56%) | 0 (0%) | 0 (0%) |
| Asian | 1 (0%) | 14 (13%) | 0 (0%) | 0 (0%) |
| Others | 23 (16%) | 1 (0%) | 0 (0%) | 0 (0%) |
| No Response | 0 (0%) | 0 (0%) | 0 (0%) | 0 (0%) |
| **Income** |  |  | N/A | N/A |
| 25-35k | 6 (4%) | 37 (34%) |  |  |
| 35-50k | 3 (2%) | 17 (16%) |  |  |
| 50-75k | 27 (19%) | 11 (10%) |  |  |
| 75-100k | 24 (17%) | 4 (4%) |  |  |
| 100-150k | 48 (34%) | 0 (0%) |  |  |
| 150-200k | 22 (15%) | 17 (16%) |  |  |
| 200k- | 12 (8%) | 12 (11%) |  |  |
| No Response | 1 (0%) | 9 (8%) |  |  |
| **Parent Education** |  |  | N/A | N/A |
| Less than college | 22 (15%) | 66 (61%) |  |  |
| College degree | 91 (64%) | 17 (16%) |  |  |
| Graduate degree | 29 (20%) | 23 (21%) |  |  |
| No Response | 1 (0%) | 2 (2%) |  |  |
| **SES Composite Score** | N/A | N/A | 3.034 ± 0.403 | 3.299 ± 0.602 |

#### Behavioral and parent-reported measures

Play and imitation behaviors were assessed using the imitation/play subscale of the Infant-Toddler Social and Emotional Assessment (ITSEA) (Briggs-Gowan & Carter, 1998; Carter et al., 2003). The subscale included six items: 1) pretends to do grown-up activities, such as shaving; 2) imitates playful sounds when prompted; 3) imitates simple adult movements, such as clapping hands or waving goodbye in response to a model; 4) pretends that objects are something else (e.g., using a banana as a phone); 5) hugs or feeds dolls or stuffed animals; and 6) rolls a ball back to another person. Parents rated each item on a three-point scale: Not True/Rarely (0 points), Somewhat True/Sometimes (1 point), or Very True/Often (2 points). The average score across these six items was used to quantify play and imitation behaviors. The timing of the brain scans relative to ITSEA assessments is illustrated in Figure S1.

Language and cognitive abilities were measured by the MacArthur-Bates Communicative Development Inventories (CDI) (Mayor & Mani, 2019) and the Mullen Scales of Early Learning (MSEL) (E. B. Lee, 2021), respectively. CDI is a widely used parent-report instrument that assesses early language development in infants and toddlers, including vocabulary comprehension, word production, and gesture use. The MSEL is a standardized developmental assessment that evaluates multiple domains of early learning. For the MSEL, we calculated the average across the five subscales: gross motor, visual reception, fine motor, receptive language, and expressive language. Autistic traits were measured with the Repetitive Behavior Scale–Revised (Wolff et al., 2016).

#### fMRI Image Acquisition

Scanning was performed at the University of North Carolina at Chapel Hill and University of Minnesota (Howell et al., 2019) on a 3T Siemens Prisma scanner with a 32-channel head coil. Structural imaging included a T1-weighted 3D-MPRAGE (TR/TE/TI = 2400/2.24/1600 ms, resolution = 0.8 mm isotropic) and a T2-weighted turbo spin-echo sequence (TR/TE = 3200/564 ms, resolution = 0.8 mm isotropic). Resting-state fMRI was acquired with a BOLD-sensitive EPI sequence (TR/TE = 800/37 ms, resolution = 2 mm isotropic, 420 volumes, 72 slices, flip angle = 80°). Additional single-band reference and AP/PA scans were obtained for distortion correction.

#### fMRI Image Processing

Preprocessed data were made available as part of the publication process by (Q. Li et al., 2024). Preprocessing was implemented using AFNI (v17.0.08) (Cox, 1996) and FSL (v6.0.1) (Jenkinson et al., 2012). Functional volumes were truncated by discarding the first 10 volumes, reoriented, and motion-corrected by alignment to the single-band EPI reference image (SBRef). Distortion correction leveraged AP/PA pairs. Functional data were co-registered with structural images (T1 for those older than 6 months; T2 for those 6 months or younger), then normalized to age-specific templates using ANTs SyN. All scans were then aligned to a common six-month template via pairwise deformation minimization. Functional data were resampled to 2 mm isotropic resolution, spatially smoothed (4 mm FWHM), detrended, and subjected to nuisance regression including 24 motion parameters, white matter, cerebrospinal fluid, global signal, and regressors for volumes with framewise displacement > 0.5 mm. Last, temporal band-pass filtering was applied (0.01–0.1 Hz).

#### fMRI Image Quality and Exclusion Criteria

A total of 292 infants with 587 task-free functional MRI scans were initially released in the BCP dataset. Data quality control followed several key steps. First, scans with incomplete volumes—fewer than the standard 420—were excluded (*N* = 8). Next, functional scans that lacked structural MRIs were excluded (*N* = 60). Scans with mean frame-wise displacement (mFD) greater than 0.5 and more than 40% high motion volumes were excluded (*N* = 42). Finally, visual inspection of all preprocessed scans led to the exclusion of 19 scans due to registration or preprocessing issues. After applying these quality control steps, the final dataset included 222 infants with 458 task-free fMRI. All scans of children under 3 years old (429 scans) were conducted during sleep, while scans of children over 3 years old (29 scans) included a mix of sleep and awake states. Of these, 143 infants had measures of play behavior.

#### Functional connectivity

Functional connectivity was calculated using preprocessed BOLD time series from each participant. Time series were extracted for all 400 Schaefer parcels, and pairwise Pearson correlations were computed between parcels to generate a 400 × 400 connectivity matrix for each participant. These correlation matrices were then Fisher z-transformed to improve normality. To facilitate network-level analyses, the parcel-wise connectivity values were subsequently averaged within each of the seven Yeo functional networks, resulting in mean connectivity estimates per network for each participant.

#### Clustering coefficient / Participation coefficient

For graph-theoretical analyses, we extracted the preprocessed time series from the same 400 Schaefer parcels. Using these data, we computed the clustering coefficient for each parcel, which quantifies the extent to which a node’s neighbors are interconnected, reflecting local network segregation. Participation coefficient was calculated for each parcel to assess the extent to which a node connects across different functional networks. For this calculation, the seven Yeo networks were used as the canonical network assignments, allowing us to quantify between-network integration. Graph metrics were computed using weighted, undirected networks derived from the Fisher z-transformed connectivity matrices. Metrics were averaged across parcels within each Yeo network to obtain network-level estimates of clustering and participation coefficients for each participant.

#### Generalized additive models

For each regional GAM, we assessed the significance of the association between the functional connectivity and age using an analysis of variance (ANOVA), comparing the full GAM model to a nested model without the age term. A significant result indicates that including a smooth term for age significantly reduced the residual deviance, as determined by the chi-squared test statistic. P-values were corrected for multiple comparisons across the six networks using the False Discovery Rate (FDR) procedure.

### Results

From ages 1 to 2 years, play scores increased sharply and then reached a plateau (Figure 2A, *R^2^* = 0.111, *P* < 0.001). Similarly, within-DMN connectivity increased until around 2 years of age (Figure 2B, *R^2^* = 0.029, *P*_FDR_ < 0.001). In contrast, within-CN connectivity remained relatively stable throughout infancy (Figure 2C, *R^2^* = 0.012, *P*_FDR_ = 0.036). Between-network connectivity generally decreased with age (Figure 2D, DMN-CN *R^2^* = 0.100, *P*_FDR_ < 0.001, Table S1), consistent with increasing functional network differentiation during early development.

**Figure 2.**
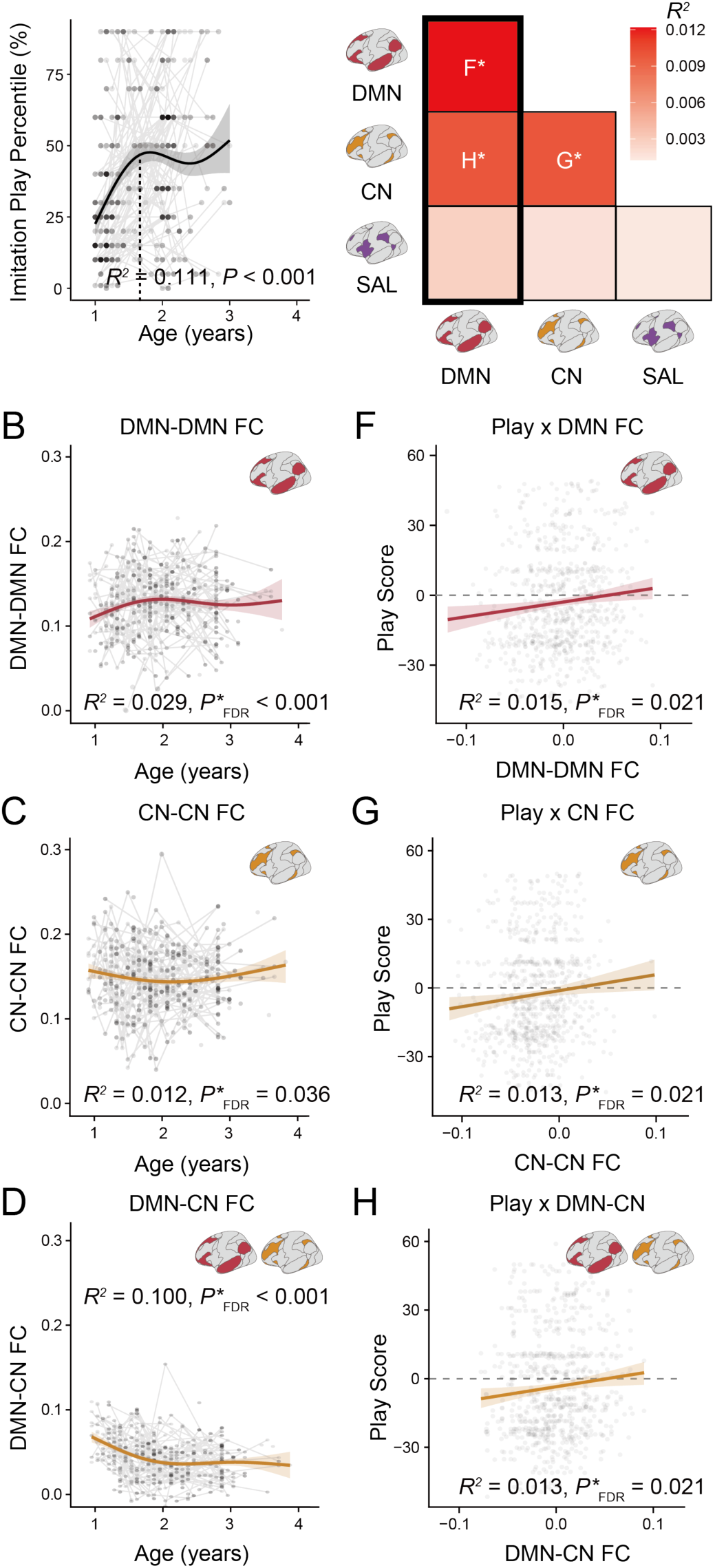
Associations between play behavior and functional connectivity. **(A)** Generalized additive model (GAM)-predicted developmental trajectory of play behavior in infancy, accompanied by a 95% confidence interval (*R^2^* = 0.111, *P* < 0.001). **(B, C, D)** GAM-predicted developmental trajectory of within DMN (B, *R^2^* = 0.029, *P*_FDR_< 0.001), within CN (C, *R^2^*= 0.012, *P*_FDR_ = 0.036) and between DMN-CN (D, *R^2^*= 0.100, *P*_FDR_ < 0.001) connectivity. **(E)** Heatmap map of correlational strength between play behavior and functional connectivity. \**P* < 0.05. **(F, G, H)** Correlation between play and within DMN (F, *R^2^* = 0.015, *P*_FDR_ = 0.021), within CN (G, *R^2^* = 0.013, *P*_FDR_ = 0.021) and between DMN-CN (H, *R^2^* = 0.013, *P*_FDR_ = 0.021) connectivity.

Controlling for age, sex and head motion, higher play scores were associated with stronger within-DMN and within-CN connectivity, and with more positive DMN-CN coupling (Figure 2E, F: Within Default: *R^2^* = 0.015, *P*_FDR_ = 0.021, Fig 2G: Within Control: *R^2^* = 0.013, *P*_FDR_ = 0.021, Fig 2H: Between Default-Control: *R^2^* = 0.013, *P*_FDR_ = 0.021). No other network pairs showed significant associations with play scores (Table S1).

We repeated our analyses controlling for potential confounds. Since opportunities for play have been previously linked to socioeconomic status (SES) (S. Li et al., 2022; Mohan & Bhat, 2022), which in turn has been linked to brain development (Kasari et al., 2006; Tooley et al., 2020), we examined whether controlling for SES altered results. In infancy, household income and parent education showed no significant relationships with play behavior (Figure S2A, B). Controlling for SES measures did not change the association between play score and brain functional connectivity (Figure S2K, L). SES was also not correlated with language or cognitive skills. We next considered whether associations between play and network connectivity were driven by cognitive abilities. Although play scores were positively correlated with language (*r* = 0.154, *P* < 0.001) and general cognitive abilities (*r* = 0.158, *P* < 0.001; Figure S2C, D), all associations between play behavior and network connectivity remained significant even after controlling for language and cognition (Figure S2M, N), suggesting that the observed brain-play relationships are at least partially independent of general cognitive development. Finally, we considered whether autistic traits influenced the play–brain association (even if the population was non-clinical), given that reduced imitation play (González-Sala et al., 2021; Kasari et al., 2006) and altered brain connectivity have been observed in individuals with autism (Padmanabhan et al., 2017). We did not find a significant correlation between autistic traits and play scores (Figure S2E), and controlling for autistic traits did not alter the brain–play associations (Figure S2O).

To assess whether the DMN, CN, and SAL emerged as key systems related to play without restricting analyses solely to predefined network-level connectivity, we additionally conducted whole-brain graph-theory analyses. The clustering coefficient indexes the extent of local connectivity among neighboring regions, reflecting the degree of local network segregation (Watts & Strogatz, 2011) (Figure 3A). The associations between clustering coefficient and age/play were generally symmetric across hemispheres; therefore, for clarity, we focus on the right hemisphere in Figure 3, and present results for the left hemisphere in Figure S3. During infancy, clustering coefficient decreased in most networks except the visual network (Figure 3B), reflecting a developmental shift toward reduced local segregation. In contrast, higher play scores were associated with higher clustering coefficients within the CN and DMN (Figure 3C), indicating greater local integration within these networks, consistent with the stronger within-network connectivity of CN and DMN reported earlier (Figure 2E-G). The participation coefficient is a graph-theoretical measure of between-network integration, indexing the extent to which a given brain region connects across multiple functional networks rather than primarily within its own network (Guimerà & Amaral, 2005) (Figure 3D). During infancy, participation coefficients increased across most networks except the visual network (Figure 3E). Play showed a modest positive association with the participation coefficients of parcels in the SAL and negative associations with participation coefficients in the CN (Figure 3F). Together, evidence from the graph metrics analysis converges with the idea that play in infancy relates to both emergent local integration within DMN and CN and selective cross-network integration involving SAL regions.

**Figure 3.**
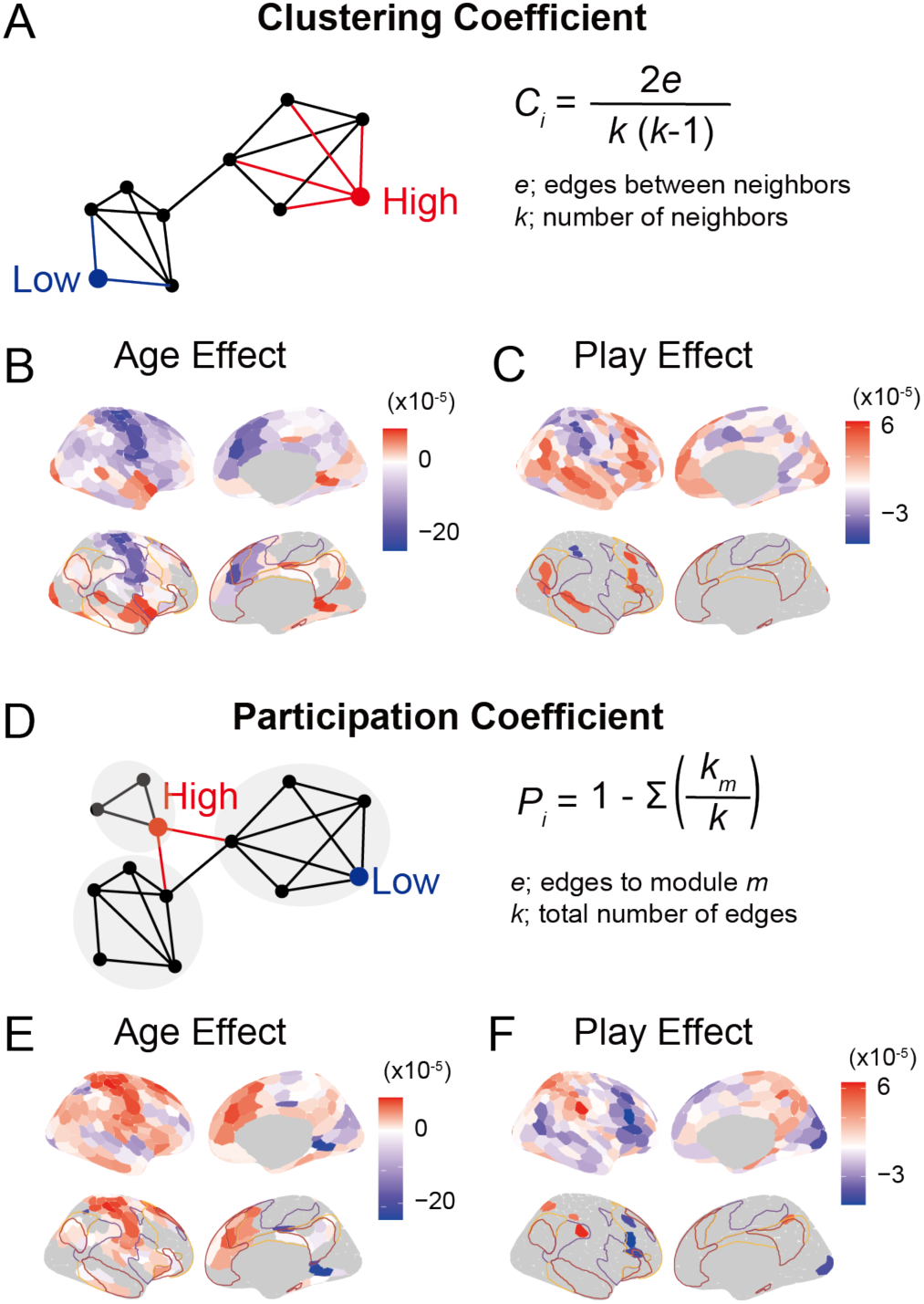
Associations between age, play and functional network measures. **(A)** Schematic of the calculation of the clustering coefficient. **(B)** Associations between age and clustering coefficients across cortex. **(C)** Associations between play and clustering coefficients across cortex. **(D)** Schematic of the calculation of the participation coefficient. **(E)** Associations between age and participation coefficients across cortex. **(F)** Associations between play and participation coefficients across cortex. For panels B,C,E and F, the upper rows show results for all brain parcels, and the lower rows display uncorrected results with *p* < 0.05. Results for the left hemisphere are presented in Figure S3.

### Discussion

Play in infancy is positively associated with stronger within- and between-network connectivity of the DMN and CN. Interestingly, during this stage, within-DMN connectivity increases and DMN-CN connectivity decreases through development, consistent with previous reports (Yin et al., 2025). Notably, these normative trends toward increasing network segregation coexist with play-related differences characterized by greater within-network coherence and stronger DMN-CN coupling, indicating that play tracks individual variability in functional integration beyond typical developmental changes.

Several limitations of this study should be acknowledged. First, the BCP collected brain imaging and play questionnaire at separate timepoints, which limited the temporal precision possible in linking neural and behavioral data. In addition, the ITSEA play questionnaire did not differentiate between types of play. Some of the items captured dramatic role-play and some captured social imitation, which may rely on different neural networks. Another limitation concerns the scanning conditions: infants in the BCP are typically scanned while asleep, which may influence the estimation of the functional connectivity. However, resting-state connectivity during sleep is thought to reflect stable, trait-like aspects of functional brain organization, and prior work has demonstrated meaningful individual differences in network architecture under these conditions. Finally, although infants’ limited ability to remain still presents substantial practical challenges, recent advances in imaging technology and motion-correction methods may help address this limitation (Ellis & Turk-Browne, 2018).

Together, and consistent with our preregistered hypotheses, Study 1 showed that play increased through infancy and was associated with stronger connectivity within the DMN, as well as greater connectivity between the DMN and CN.

## Study 2

Building on Study 1, which examined the emergence of play in infancy, Study 2 investigates whether age-related changes in connectivity among the DMN, CN, and SAL correspond to the decline of play behaviors during middle childhood.

### Method

This study was not preregistered.

#### Participants

The childhood dataset was collected at the University of Pennsylvania with Institutional Review Board approval. A subset of these data were previously published (*N* = 92) (Tooley et al., 2022). Children were recruited from Philadelphia and surrounding areas through schools, advertisements, outreach programs, and community events. Exclusion criteria included diagnosis of neurological, mental or developmental disorder, adoption, and MRI contraindications. Parents provided informed written consent; children gave verbal assent (if < 8 years old) or written assent (≥ 8 years old). Participants ranged from 4.11–10.59 years (M = 6.55, SD = 1.36; 47% male, 53% female). Demographics are shown in Table 1. The final sample included 128 children after exclusions for incomplete resting-state scans (*N* = 17), undisclosed diagnoses based on Child Behavior Checklist scores (*N* = 4), and additional data quality criteria described below (*N* = 40). The participants’ demographic characteristics are shown in Table 1.

#### Parent-reported measures

Play behavior was measured using items from the Home Literacy and Numeracy questionnaire (Manolitsis et al., 2013). The play frequency section consists of twelve items assessing the frequency of engagement in activities such as: 1) Sorting things by size, color, or shape, 2) Putting pegs in a board or shapes into holes, playing with puzzles; 3) Building Lego or construction set (Duplo, Megablocks, etc.); 4) Playing with “Playdoh” or clay; 5) Playing with blocks; 6) Coloring, painting, writing; 7) Doing crafts involving scissors and glue; 8) “Paint-by-number” activities; 9) Movement songs (i.e., Itsy Bitsy Spider); 10) Playing musical instruments; 11) Playing store and 12) Playing teacher.

Responses were recorded on a six-point scale: Did not occur, Less than once a week but a few times in a month, Once a week, A few times a week, Almost daily, Activity not applicable to your child, or Prefer not to answer.

#### fMRI Image Acquisition

Scanning was conducted at the University of Pennsylvania on a Siemens MAGNETOM Prisma 3T scanner with a 32-channel head coil. Structural imaging included a T1-weighted multi-echo MEMPRAGE (TR = 2530 ms, TI = 1330 ms, voxel size = 1 mm isotropic). Resting-state fMRI was acquired with a BOLD-sensitive sequence (TR = 2000 ms, TE = 30.2 ms, resolution = 2 mm isotropic, flip angle = 90°). Children underwent a mock scan prior to imaging and were monitored with the Framewise Integrated Real-time MRI Monitor system. When possible, 10 minutes of low-motion resting-state data (two runs, FD < 1 mm) were collected.

#### fMRI Image Processing

Preprocessing was implemented using fMRIPrep (v20.2.0) and XCP-D (v0.5.0) (Ciric et al., 2018), with ANTs, FSL (Jenkinson et al., 2002), and AFNI (Cox, 1996). The T1-weighted image underwent intensity nonuniformity correction, skull stripping, and spatial normalization to the ICBM 152 Nonlinear 2009c atlas, followed by tissue segmentation into CSF, white matter, and gray matter. For each resting-state BOLD run, a reference volume was generated, BOLD data were coregistered to the T1-weighted reference using boundary-based registration, and head-motion parameters were estimated using *mcflirt*. Slice timing correction was applied, and BOLD time series were resampled to MNIPediatricAsym space using a single composite transform with Lanczos interpolation. Confounding time series including framewise displacement, DVARS, and global signals were calculated, and nuisance regression using a 36-parameter model was applied on the fsLR cortical surface. Motion censoring flagged volumes exceeding FD = 0.3 mm, which were interpolated using cubic splines, and runs with less than 100 seconds of usable data were excluded.

#### fMRI Image Quality and Exclusion Criteria

The imaging data quality was evaluated through fMRIPrep visual reports and MRIQC 0.14.2 software. Two reviewers manually evaluated all structural and functional images at each preprocessing stage for any image quality concerns. Functional images underwent visual inspection to ensure adequate whole-brain field of view coverage, absence of signal blurring or artifacts, and correct alignment with the anatomical image. Exclusions were made for participants with unusable structural images (*N* = 1), functional data artifacts (e.g., hair glitter, *N* = 1), incorrect scanner registration (*N* = 1), registration failure (*N* = 18), less than 100 seconds usable data after high-motion outlier volumes (*N* = 13), and average framewise displacement > 1 mm (*N* = 6). For participants with multiple usable resting-state runs, framewise displacement was averaged across runs.

#### Network Analyses

Functional connectivity, clustering coefficients, and participation coefficients were calculated in the same was as in Study 1. Generalized additive models were used as in Study 1.

### Results

Between ages 4 and 12 years old, parent-reported play frequency declined linearly (Figure 4A *R^2^* = 0.042, *P* = 0.035). In parallel, DMN–SAL and CN–SAL connectivity also decreased linearly throughout childhood (Figure 4B *R^2^* = 0.074, *P*_FDR_ = 0.007, C *R^2^* = 0.008, *P*_FDR_ = 0.366), consistent with increasing large-scale network segregation during childhood. Higher play scores were associated with stronger DMN–SAL and CN–SAL connectivity (Figure 4D, E *R^2^* = 0.080, *P*_FDR_ = 0.012, F *R^2^* = 0.067, *P*_FDR_ = 0.012), suggesting that children who engaged more in play were characterized by greater cross-network integration, particularly involving the SAL. No other network pairs showed significant associations with play scores (Table S2).

**Figure 4.**
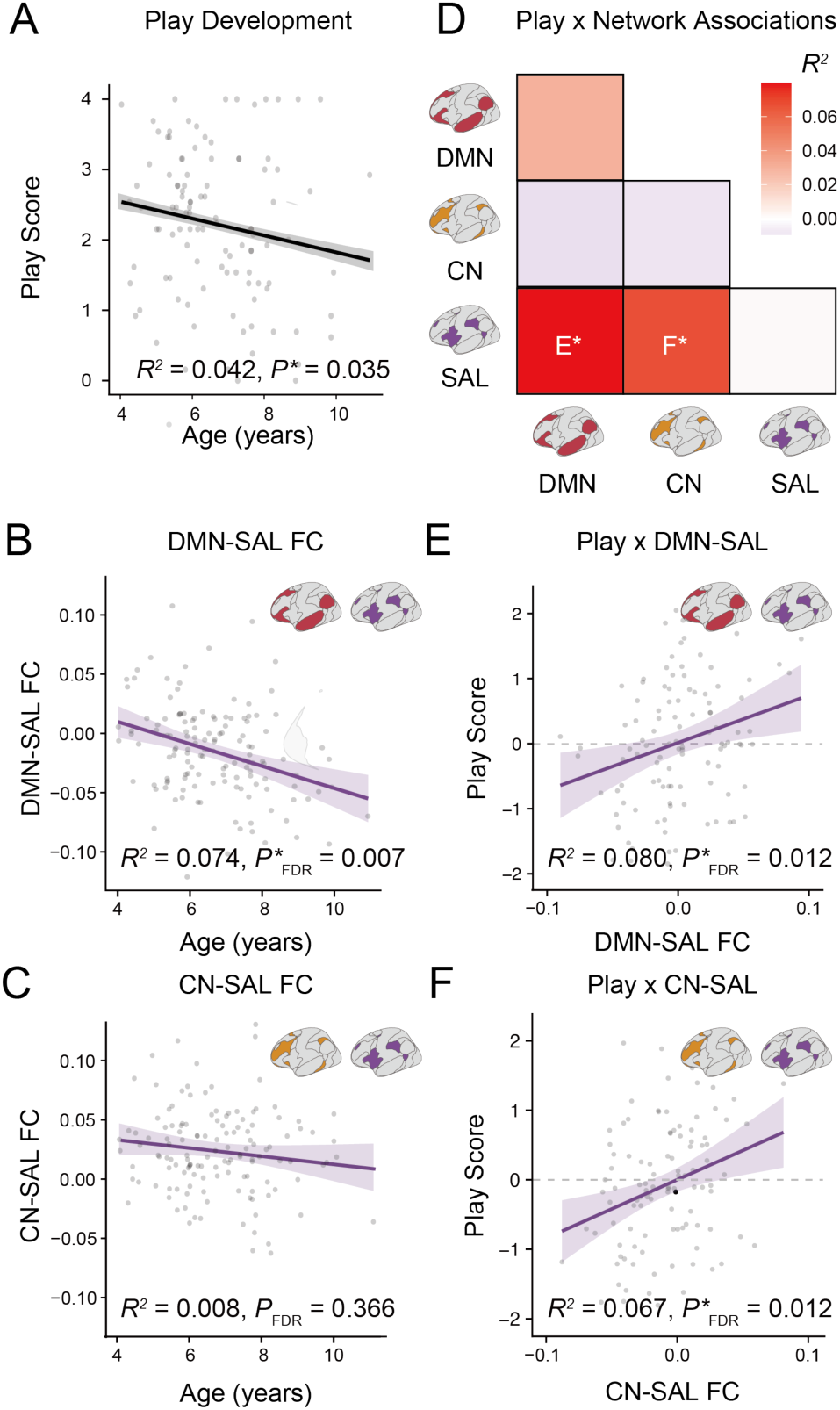
Associations between play behavior and functional connectivity. **(A)** Generalized additive model (GAM) predicted developmental trajectory of play behavior in childhood, accompanied by a 95% confidence interval (*R^2^* = 0.042, *P* = 0.035). **(B, C)** GAM-predicted developmental trajectory of between DMN-SAL (B, *R^2^* = 0.074, *P*_FDR_ = 0.007) and between CN-SAL (C, *R^2^* = 0.008, *P*_FDR_ = 0.366) connectivity. **(D)** Heatmap map of correlational strength between play behavior and functional connectivity. \**P* < 0.05. **(E, F)** Correlation between play and between DMN-SAL (E, *R^2^* = 0.080, *P*_FDR_ = 0.012) and between CN-SAL (F, *R^2^* = 0.067, *P*_FDR_ = 0.012) connectivity. vb

Household income and parental education were negatively associated with play frequency (Figure S4A,B), a pattern that contrasts with some prior reports and motivated sensitivity analyses controlling for socioeconomic factors. Controlling for median income or parental education, the DMN-SAL association with play was attenuated and no longer survived FDR correction (Figure S4C; *R*^2^ = 0.039, *P* = 0.049, *P*_FDR_ = 0.093, D; *R*^2^ = 0.032, *P* = 0.038, *P*_FDR_ = 0.113), whereas CN-SAL remained robust (Figure S4E *R*^2^ = 0.065, *P* = 0.007, *P*_FDR_ = 0.045, F; *R*^2^ = 0.071, *P* = 0.004, *P*_FDR_ = 0.022).

Across childhood, clustering coefficients generally showed weak negative associations with age in lateral prefrontal and superior temporal areas and showed significantly positive associations with age in visual areas (Figure 5A). None of the clustering coefficients for any brain parcel showed a significant association with play (Figure 5B), indicating that play-related effects in childhood were not driven by differences in local network segregation. Participation coefficients showed broad negative associations with age, with significant effects in visual and medial prefrontal areas (Figure 5C). A few parcels within the DMN, CN, and SAL showed positive associations with play (Figure 5D), although these parcel-level effects were sparse and did not survive correction for multiple comparisons, consistent with the positive relationship between play and between-network functional connectivity involving the SAL (Figure 4D-F).

**Figure 5.**
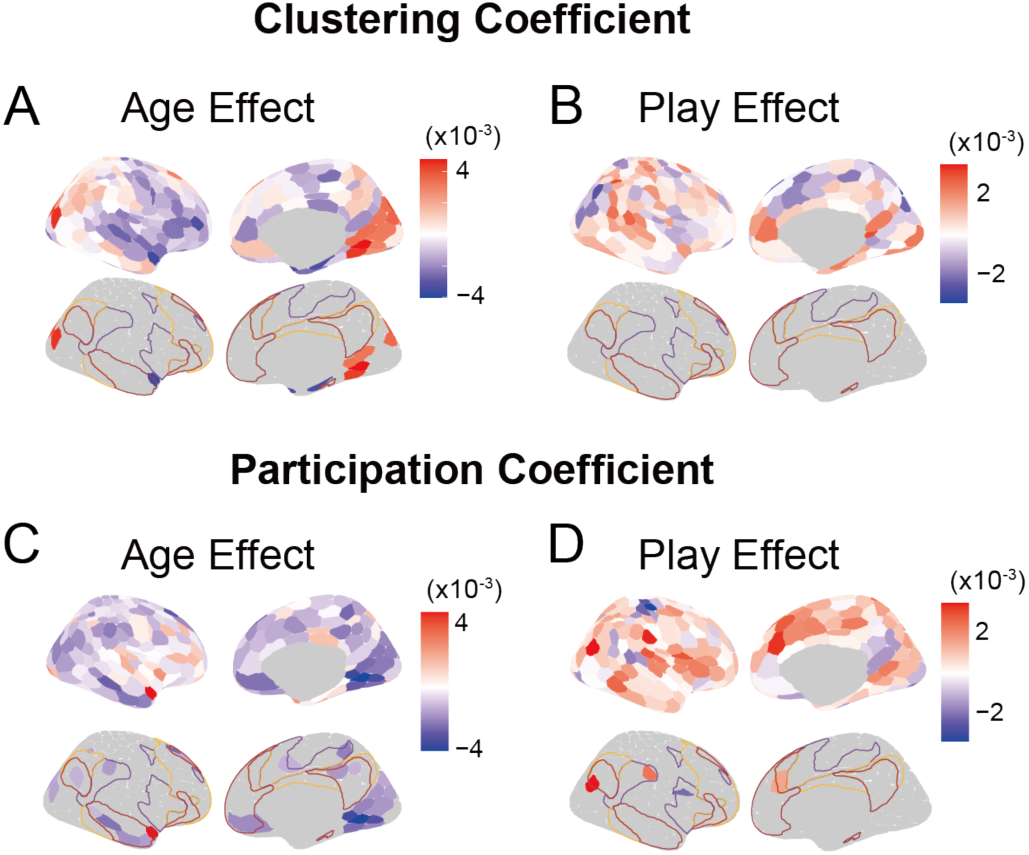
Associations between age, play and functional network measures. **(A)** Associations between age and clustering coefficients across cortex. **(B)** Associations between play and clustering coefficients across cortex. **(C)** Associations between age participation coefficients across cortex. **(D)** Associations between play and participation coefficients across cortex. For panels A-D, the upper rows show results for all brain parcels, and the lower rows display uncorrected results with *p* < 0.05. Results for the left hemisphere are presented in Figure S5.

### Discussion

Children who engaged in more play showed relatively greater connectivity between DMN-SAL and CN-SAL networks, suggesting that play may help preserve network integration. Overall, DMN-SAL and CN-SAL connectivity decreased through childhood, consistent with previous reports of a general trend toward increasing network segregation (Tooley et al., 2022).

One limitation to note is that the Home Literacy and Numeracy questionnaire (Manolitsis et al., 2013) does not differentiate between types of play; it includes sensorimotor play as well as symbolic and dramatic pretend play, which may depend on different forms of neural network integration. This heterogeneity may contribute to the prominence of salience-related effects in childhood, given the SAL’s role in coordinating diverse cognitive and behavioral demands. Moreover, the questionnaire primarily captures free, unstructured play, while structured activities, including guided play, playful learning, or more social forms of play that remain important in childhood, are not well represented. Therefore, the decline in play scores observed in this study may not necessarily reflect a true decrease in play behavior. It is also possible that it reflects a transition or change in the forms of play (Hofferth & Sandberg, 2001; Weisberg et al., 2016; Zosh et al., 2018), which our questionnaire was unable to capture.

In summary, Study 2 showed that higher levels of play were associated with preserved integration among large-scale brain networks in childhood, specifically greater between-network connectivity between DMN-SAL, as well as between CN-SAL. Notably, this pattern resembles that seen in infancy (Study 1), where play was most strongly associated with integration between the DMN and CN, highlighting a developmental reorganization in the network systems linked to play across development.

## Study 3

While the results of Study 1 and 2 provide converging evidence for an association between play behavior and functional brain network integration development, they do not resolve the directionality of this relationship—specifically, whether children with more integrated brain networks are more likely to engage in play, or whether children educated in Montessori settings, learning environments that emphasize child-led, materials-rich, and hands-on play-like exploration, exhibit greater network integration among DMN, CN, and SAL compare to peers in traditional schools.

### Method

This study was not preregistered.

#### Participants

The Montessori dataset was previously collected and published by (Duval et al., 2023). Participants were drawn from over 20 schools across Switzerland, representing either Montessori or traditional educational settings. Eligibility required children to be between 4 and 18 years old and to have a consistent educational background (Montessori or traditional for at least three years, or since age 4 for children younger than 7 years old). Children were excluded if parents reported learning or behavioral difficulties, or if they had sensory impairments. Since Montessori schools in Switzerland are exclusively private, recruitment was geographically restricted to regions with traditional schools matched for higher-income levels, based on official city salary statistics. Ethical approval was granted by the Commission d’Ethique Romande Vaud (CER-VD 2018-00244), and all procedures followed the Declaration of Helsinki. Written informed consent was obtained from parents, with verbal assent from the children.

#### SES composite score

An SES composite score was derived from a parental questionnaire (Genoud, 2011) that gathered information on professional status, occupational category, and education level. Higher scores indicated higher SES (*M* = 3.117, *SD* = 0.532, range 2-4). After narrowing the age range to under 12 years to align with the Study 2 dataset and applying further data quality checks (described below), the final sample comprised 44 children. The participants’ demographic characteristics are shown in Table 1. The groups did not significantly differ by age or sex, but the children in the traditional schooling group did come from slightly higher SES backgrounds (Figure S7A, *t* = 1.742, *p* = 0.089).

#### fMRI Image Acquisition

Scanning was performed at the CIBM Center for Biomedical Imaging at Lausanne University Hospital on a Siemens 3T Prisma-Fit scanner with a 64-channel head coil. Structural imaging included a T1-weighted scan (TR/TE/TI = 2000/2.47/900 ms, flip angle = 8°, voxel size = 1 mm isotropic, FOV = 256 × 256 mm²). Resting-state fMRI was acquired with a BOLD-sensitive EPI sequence (TR/TE = 500/33 ms, flip angle = 80°, voxel size = 2.4 mm isotropic, matrix size = 94 × 94, FOV = 224 × 224 mm², 48 slices, 720 volumes, scan duration = 6 min, slice acceleration factor = 8, bandwidth = 2418 Hz/Px).

#### fMRI Image Processing

The preprocessing was implemented using fMRIPrep (v20.2.1) and XCP-D (v0.12.0) (Ciric et al., 2018). The Montessori pipeline applied band-pass filtering between 0.01–0.08 Hz using a second-order Butterworth filter and included a despiking step, and no frame censoring or motion filtering was applied.

#### fMRI Image Quality and Exclusion Criteria

Six reviewers manually evaluated all structural and functional images and rated their quality as poor (dental brace interferences, FD and DVARS <1.5), medium (FD and DVARS mean values in range, but peaks outside range), or good. Participants with poor or medium image quality were excluded (*N* = 10).

#### Network Analysis

We examined the relationship between brain network connectivity and school environments using linear regression. Both the network measures and school environments were residualized for covariates (age, sex, motion, and, in the sensitivity analysis, the SES composite score) to isolate variance independent of these factors. Each residualized network was regressed onto the residualized outcome, and the resulting p-values were corrected for multiple comparisons across the six networks using the FDR procedure.

### Results

Children in Montessori schools showed higher DMN-SAL connectivity (Figure 6B, *β* = 3.48, *P*_FDR_ = 0.049), as well as DMN-CN connectivity (Figure 6C, *β* = 3.55, *P*_FDR_ = 0.049) than those in traditional schools. No other significant differences were observed. No school environment by age interactions were observed (Table S3, Figure S6). While the SES composite scores did not differ significantly between the school environments, SES was slightly higher in children attending traditional schools. When controlling for SES composite score, the differences between school environments for both DMN–SAL and DMN–CN connectivity were attenuated and no longer survived FDR correction (Figure S7C, DMN-SAL *t* = 2.939, *P* = 0.056, *P*_FDR_ = 0.127, DMN-CN *t* = 3.087, *P* = 0.020, *P*_FDR_ = 0.120).

**Figure 6.**
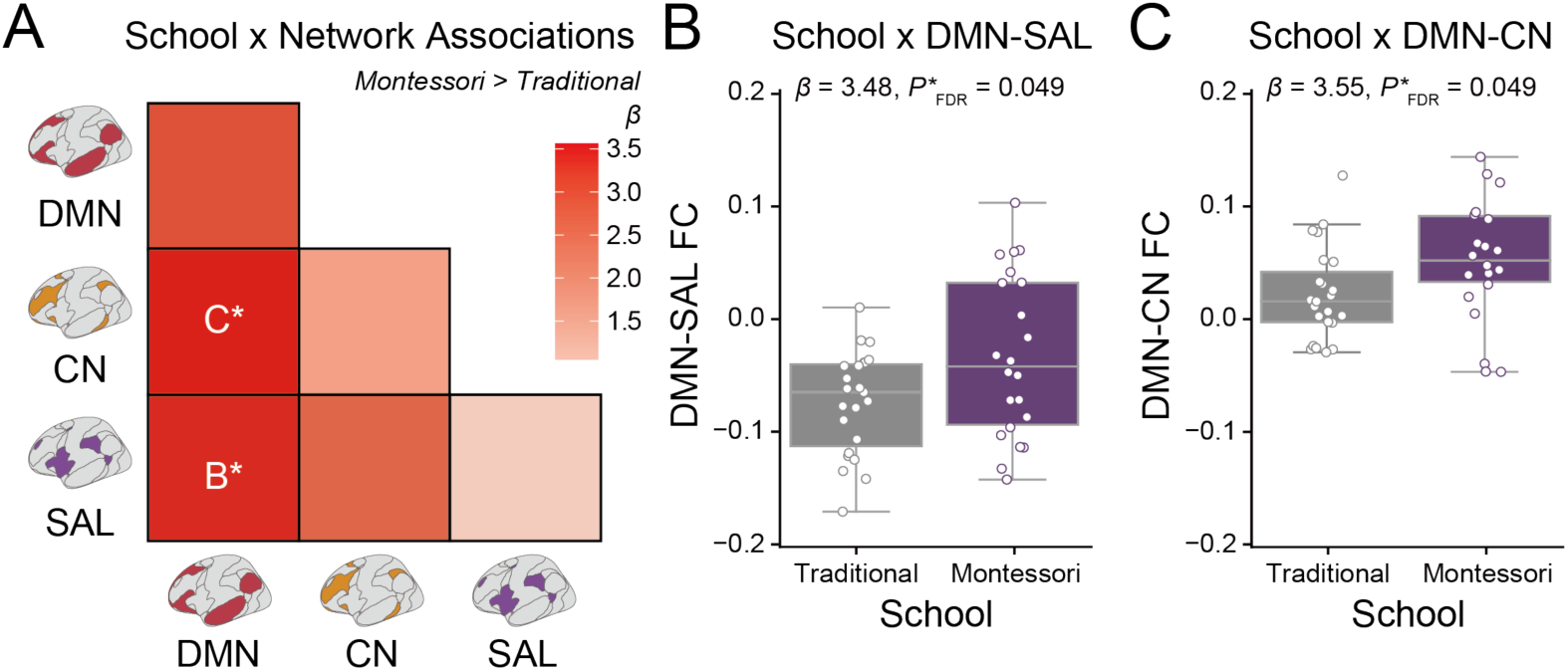
Relationship between school environment and brain functional network connectivity. **(A)** Heatmap showing the strength of correlations between play behavior and functional connectivity. \**P*_FDR_ < 0.05. Age, sex, and mean framewise displacement were controlled as covariates. **(B)** Group comparison of DMN-SAL connectivity between children in traditional (gray) and Montessori (purple) schools (*β* = 3.48, *P*_FDR_ = 0.049). **(C)** Group comparison of DMN-CN connectivity between children in traditional (gray) and Montessori (purple) schools (*β* = 3.55, *P*_FDR_ = 0.049).

### Discussion

Children who attended Montessori programs showed higher DMN–SAL and DMN–CN functional connectivity than peers in traditional schools, suggesting that environments which facilitate playful learning may help maintain or enhance functional integrations. These findings are consistent with the broader pattern observed in Study 1 and 2, in which play and play-like exploration were associated with greater integration among large-scale brain networks. Importantly, however, the attenuation of effects after controlling for SES indicates that school-related differences in network integration cannot be attributed uniquely to educational context.

Because Study 3 did not include direct measures of play, it remains unclear whether differences in play opportunities contributed to the observed differences between school environments, or whether other features of the Montessori approach, such as child-led exploration, multisensory learning, or minimal external evaluation (Denervaud et al., 2019; Denervaud, Fornari, et al., 2020; Denervaud, Knebel, et al., 2020; Denervaud, Mumenthaler, et al., 2020; Lopata et al., 2005; Rathunde & Csikszentmihalyi, 2005), played a larger role. In addition, because children were not randomly assigned to school type, these findings are correlational and may reflect selection effects, such as pre-existing differences in cognitive style, family values, or socioeconomic factors, rather than causal effects of the educational environment itself.

Overall, Study 3 provides convergent but qualified evidence that educational environments that promote play-like exploration, such as Montessori programs, are associated with greater functional integration among large-scale brain networks, although causal mechanisms remain to be established.

## General Discussion

Across three studies spanning infancy to middle childhood, we found converging evidence that play is associated with greater integration both within and between large-scale brain networks. In infancy, as play behavior increases rapidly, more frequent play was associated with stronger within-network connectivity for the DMN and CN, and greater DMN-CN coupling. During childhood, when play behavior declines, more frequent play was associated with preserved integration of SAL with DMN and with CN. Moreover, children in Montessori schools showed stronger connectivity between DMN-CN and DMN-SAL than peers in traditional schools, suggesting that learning environments centering child-led play may support the integration of the brain networks. Together, these findings are consistent with our pre-registered hypothesis that more frequent play is associated with greater integration between the DMN and other large-scale functional networks, a pattern previously linked to creative cognition in adults (Beaty et al., 2014, 2016, 2018; Hoffmann & Russ, 2012; S. Lee et al., 2021).

We consistently observed that more frequent play is associated with greater integration of the DMN, CN, and SAL in both infancy (Study 1) and childhood (Study 2), a pattern also seen in highly creative adults (Beaty et al., 2014, 2018). Although play is not always goal-directed, children often generate functional goals (e.g. “build a tower,” “connect the puzzle pieces”) and imaginative goals (e.g., “save the princess,” “host a feast”) and monitor progress toward them, thereby exercising flexible modulation between more integrated and more segregated network states as they shift between ideation and evaluation (Jung et al., 2013). This dynamic may support both the development and coordination of DMN and CN, as well as the ability to flexibly switch between them. In addition, by linking external stimuli and objects to imaginative scenarios (such as a cardboard box as a spaceship or a puddle as a magical portal), play may also enhance coordination between the DMN and SAL. While the increasing segregation of brain networks is a hallmark of typical childhood development (He et al., 2019; Tooley et al., 2022) and is essential for efficient information processing and task performance (Sims et al., 2022; R. Wang et al., 2021), this segregation may, in some contexts, be associated with reduced creativity and imagination.

The greater integration of DMN, CN and SAL observed in children attending Montessori schools in Study 3 is consistent with these ideas. Montessori emphasizes child-led exploration, multisensory materials, minimal extrinsic evaluation, and no time pressure (Rand & Montessori, 1941; Sobe, 2004). Although these features overlap with key characteristics of play, such as autonomy, exploration, and intrinsic motivation, they are not synonymous with play per se and may engage partially distinct cognitive processes. In contrast, traditional schooling typically places greater emphasis on teacher-directed instruction, structured curricula, and external evaluation, though practices vary widely across educational contexts. Prior research has shown that Montessori students often demonstrate greater creative thinking, knowledge integration, and self-monitoring compared to children attending traditional schools (Denervaud et al., 2019; Denervaud, Fornari, et al., 2020; Denervaud, Knebel, et al., 2020; Denervaud, Mumenthaler, et al., 2020; Lopata et al., 2005; Rathunde & Csikszentmihalyi, 2005). Taken together with the present findings, it is possible that more frequent opportunities for learning through play in Montessori education foster integration of the DMN, CN and SAL, which in turn could contribute to higher creativity (Beaty et al., 2018; Denervaud et al., 2019; Denervaud, Fornari, et al., 2020; Denervaud, Knebel, et al., 2020; Denervaud, Mumenthaler, et al., 2020; Duval et al., 2023).

Although previous studies have reported a negative association between SES and play in infancy (Clearfield et al., 2014; S. Li et al., 2022; Mohan & Bhat, 2022), we did not find any significant relationship between SES and play in Study 1. This could be due to limited variation in SES within the dataset, as over 90% of parents in the dataset hold a college degree or higher. The lack of correlation between SES and language or cognitive skills further supports this explanation. In Study 2, a negative correlation between SES and play was observed, suggesting that children from lower-SES households engaged in play more frequently. This negative correlation may suggest several possibilities. One is that parents in high-SES households are more likely to be dual-income earners, so their children are watched by other caregivers during the day, which may limit their opportunities to observe their children engaging in play. Another possibility is that high-SES families may place greater emphasis on academic achievement (Z. Li & Qiu, 2018) or encourage participation in extracurricular activities, which may reduce opportunities for play. Speculatively, several non-mutually exclusive factors may contribute to this relationship, including differences in caregiving arrangements, parental expectations, or the structure of children’s daily activities across socioeconomic contexts.

Several limitations of this work should be acknowledged. Importantly, these limitations also highlight key directions for future research aimed at establishing causal links between play behavior and brain network development. First, our measures of play relied on parent reports and included only a few items for each type of play, which limits the precision and depth of the behavioral data. In addition, most of these measures focus on play conducted by children alone and therefore do not fully capture important social aspects, such as interactions with parents and other children. Direct observational assessments of play, both at home and in school, with differentiation between play types (e.g., sensorimotor, symbolic, sociodramatic) and inclusion of social dynamics, such as dyadic interactions, would provide a more detailed and accurate characterization of play-related behaviors. Second, all data were correlational, precluding causal inference, although Study 3 provides quasi-experimental evidence through comparison of educational contexts, school assignment was non-random and remains subject to selection effects. Future studies could strengthen causal claims through longitudinal designs, experimental interventions, or assignment to play-promoting environments, such as Montessori programs, to test whether changes in play lead to measurable changes in brain network development. Additionally, the sample sizes across studies are relatively small (*N* = 143 for Study 1, *N* = 108 for Study 2, *N* = 44 for Study 3) and the observed effect sizes were generally modest; however, similar patterns emerged across independent samples, age ranges, and analytic approaches, supporting the robustness of the core associations. Parcel-level analyses did not reveal any effects that survived FDR correction, suggesting that the relationship between play and cortical connectivity is not localized to specific brain regions and is likely subtle. Furthermore, our analyses focused on cortical networks defined using an adult atlas (Thomas Yeo et al., 2011), which may be less accurate for infants, and did not include subcortical contributions, such as thalamic or hippocampal connectivity (Sydnor, 2025). Future work incorporating infancy- and childhood-specific functional atlases (Tooley, Bassett, et al., 2022; F. Wang et al., 2023) and explicitly examining subcortical dynamics could further clarify the neural substrates of play.

Across three studies, more frequent play was consistently associated with greater functional integration among large-scale brain networks, including the DMN, CN, and SAL, with a developmental shift from play-related integration primarily between the DMN and CN in infancy to SN-mediated integration in childhood. As academic pressures increase and young children gain earlier and easier access to virtual activities and digital games, opportunities for play may be diminishing in some contexts. The present findings generate the hypothesis that such shifts could involve neural trade-offs of reduced play: prioritization of segregation beneficial for focused performance versus integration that may scaffold creativity and adaptive switching, skills crucial for children’s future in a tech-driven era.

## Transparency and Openness

Data for Study 1 were obtained from the Baby Connectome Project (BCP), publicly available via the NIMH Data Archive (NDA; https://nda.nih.gov/), with access requiring an approved Data Use Certification. The derivative data needed to replicate Studies 1–3 are provided with the analysis code, which is publicly available on GitHub at https://github.com/monami-nishio/play_functional_connectivity.

