## Supplemental Materials for "Developmental links between play behavior and brain network integration"

**Online Supplemental Materials**

Developmental links between play behavior

and brain network integration

**Date of submission:** March, 3, 2026.

### Supplementary Materials: Study 1

**Supplementary Table 1**

*Associations between network connectivity and age/play in infancy*

|  | Age | | | Play | | |
| --- | --- | --- | --- | --- | --- | --- |
|  | *R^2^*_Partial_ | *P* | *P_FDR_* | *R^2^*_Partial_ | *P* | *P_FDR_* |
| DMN-DMN | 0.029 | <0.001 | <0.001 | 0.015 | 0.004 | 0.021 |
| DMN-CN | 0.100 | <0.001 | <0.001 | 0.013 | 0.011 | 0.021 |
| DMN-SAL | 0.024 | <0.001 | <0.001 | 0.005 | 0.270 | 0.405 |
| CN-CN | 0.012 | 0.030 | 0.036 | 0.013 | 0.008 | 0.021 |
| CN-SAL | 0.028 | <0.001 | <0.001 | 0.004 | 0.359 | 0.431 |
| SAL-SAL | 0.005 | 0.171 | 0.171 | 0.003 | 0.953 | 0.953 |

**Supplementary Figure 1**

***Timing of scans and questionnaires for each participant***

The timing of the scan (red, circle) and the play questionnaire (blue, cross) is shown for each participant. Participants are ordered by the timing of their earliest scan.

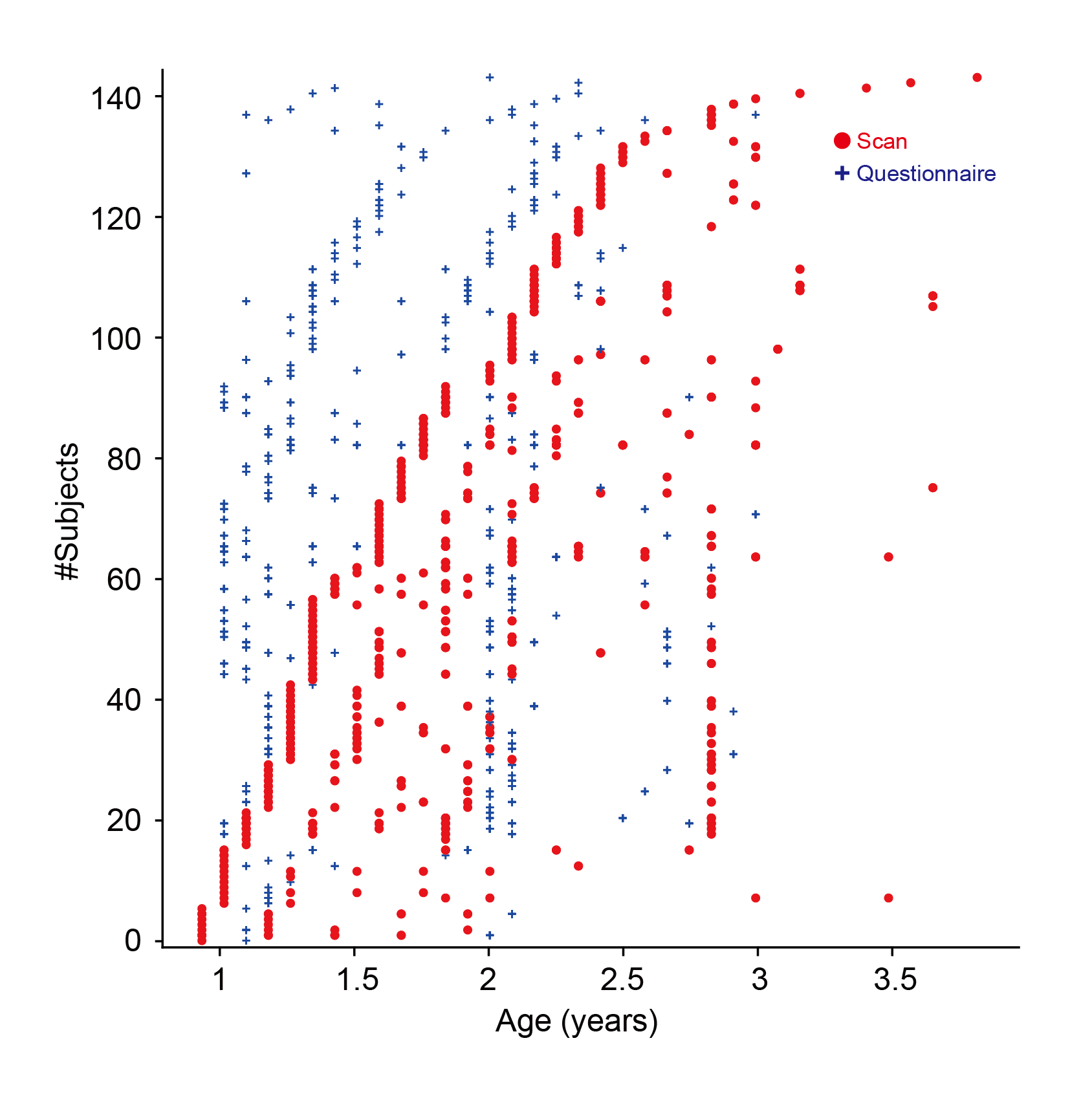

**Supplementary Figure 2. Sensitivity analysis. (A-E)** Correlations between play behavior and household income (A, *r* = 0.025, *P* = 0.681), parent education (B, *r* = -0.006, *P* = 0.930), language score (C, *r* = 0.154, *P* < 0.001), cognitive score (D, *r* = 0.158, *P* < 0.001) and autism spectrum disorder (ASD) score (E, *r* = -0.031, *P* = 0.444). **(F-J)** Heatmap map of correlational strength between functional connectivity and household income (F), parent education (G), language score (H), cognitive score (I) and ASD score (J). **(K-O)** Heatmap map of correlational strength between functional connectivity and play behavior controlling for household income (K), parent education (L), language score (M), cognitive score (N) and ASD score (O). **P* < 0.05.

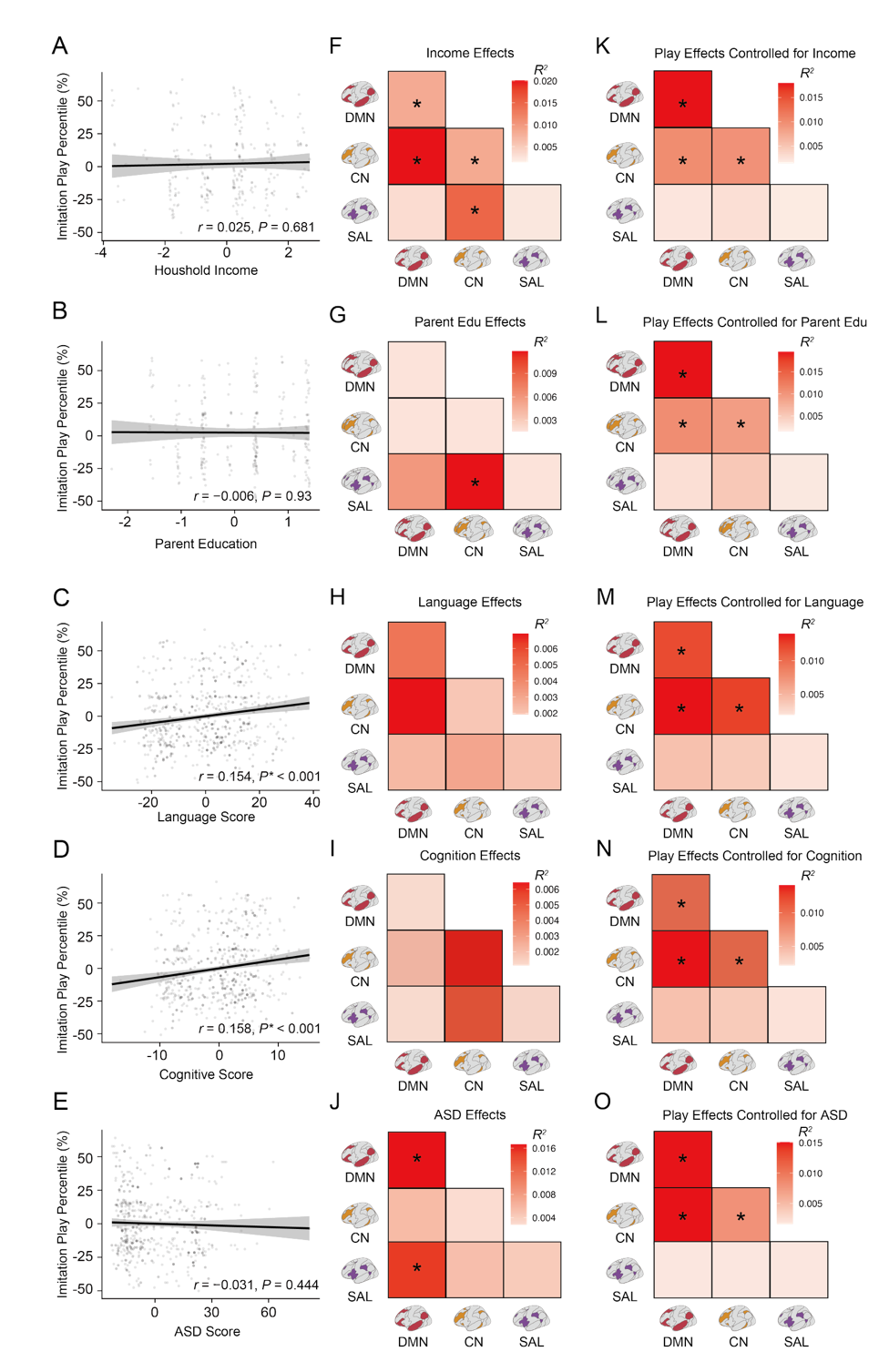

**Supplementary Figure 2**

*Sensitivity analysis*

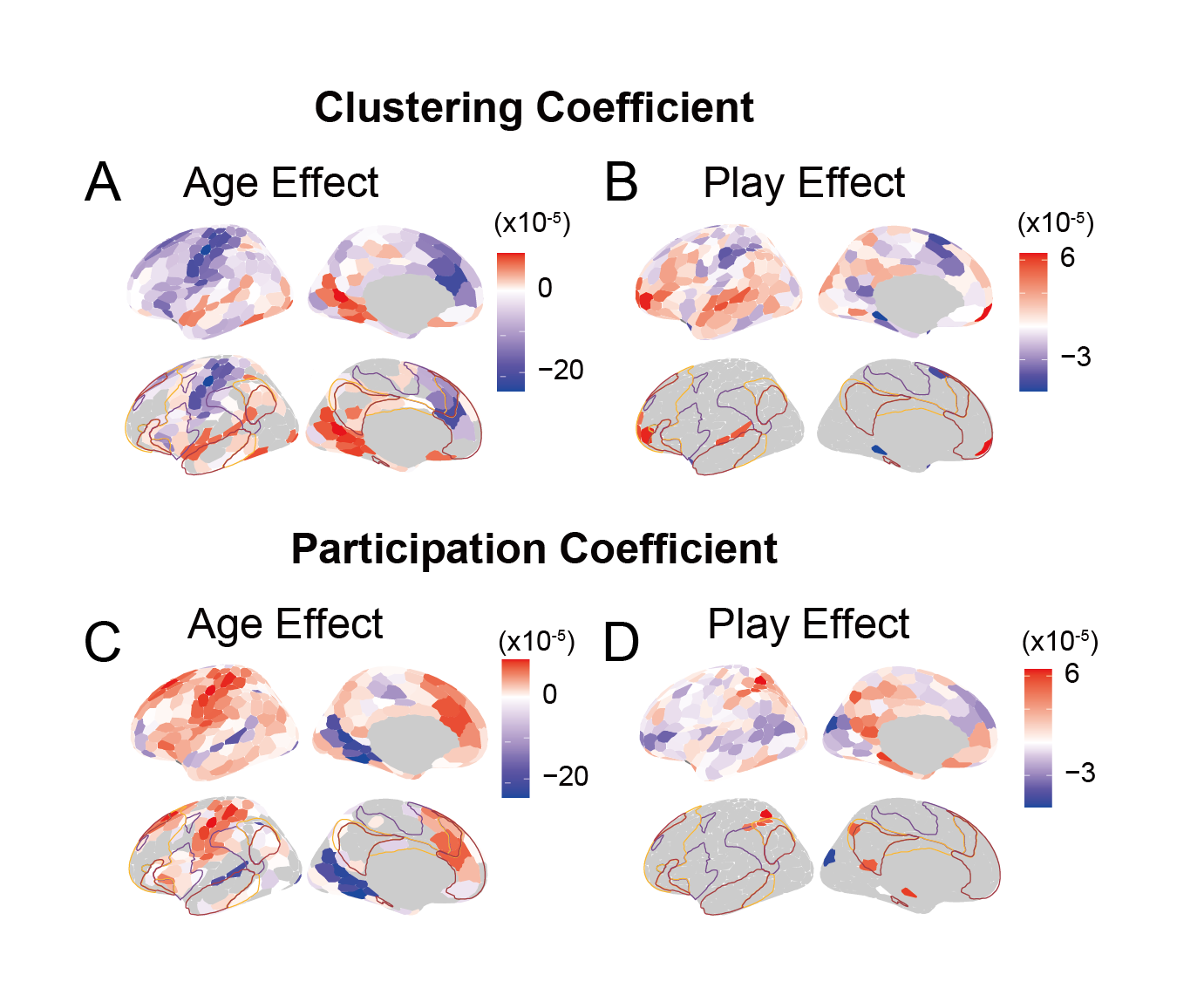

**Supplementary Figure 3. Associations between age, play and functional network measures in the left hemisphere in infancy. (A)** Associations between age and clustering coefficients across cortex. **(B)** Associations between play and clustering coefficients across cortex. **(C)** Associations between age and participation coefficients across cortex. **(D)** Associations between play and participation coefficients across cortex. For panels A-D, the upper rows show results for all brain parcels, and the lower rows display uncorrected results with *p* < 0.05.

**Supplementary Figure 3**

*Associations between age, play and functional network measures in the left hemisphere in infancy*

### Supplementary Materials: Study 2

|  | Age | | | Play | | |
| --- | --- | --- | --- | --- | --- | --- |
|  | *R^2^* _Partial_ | *P* | *P_FDR_* | *R^2^* _Partial_ | *P* | *P_FDR_* |
| DMN-DMN | 0.044 | 0.013 | 0.027 | 0.030 | 0.042 | 0.084 |
| DMN-CN | 0.006 | 0.389 | 0.389 | -0.010 | 0.933 | 0.933 |
| DMN-SAL | 0.074 | 0.001 | 0.007 | 0.080 | 0.002 | 0.012 |
| CN-CN | 0.032 | 0.038 | 0.056 | -0.007 | 0.625 | 0.750 |
| CN-SAL | 0.008 | 0.305 | 0.366 | 0.067 | 0.004 | 0.012 |
| SAL-SAL | 0.058 | 0.004 | 0.013 | 0.002 | 0.281 | 0.422 |

**Supplementary Table 2**

*Associations between network connectivity and age/play in childhood*

**Supplementary Figure 4**

*Sensitivity analysis*

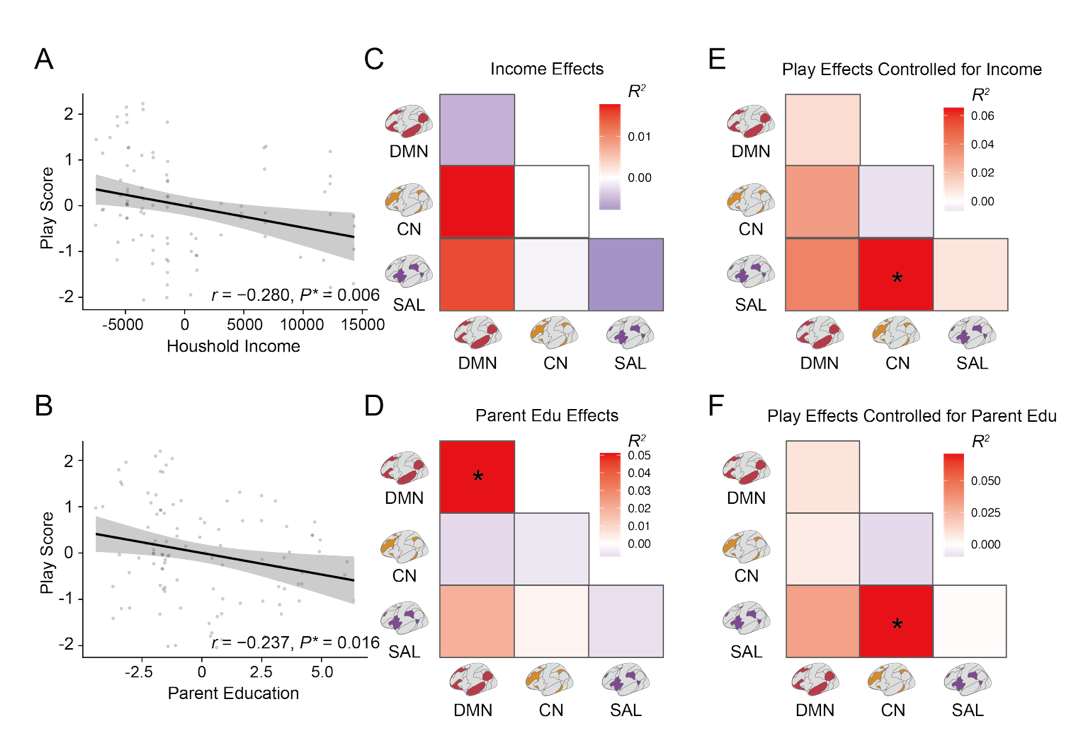

**(A, B)** Correlation between play behavior and household income (A, *r* = -0.280, *P* = 0.006) and parent education (B, *r* = -0.237, *P* = 0.016) in childhood.  **(C, D)** Heatmap map of correlational strength between functional connectivity and household income (C) and parent education (D) in childhood. **(E, F)** Heatmap map of correlational strength between functional connectivity and play behavior controlling for household income (E), and parent education (F).

**(A)** Associations between age and clustering coefficients across cortex. **(B)** Associations between play and clustering coefficients across cortex. **(C)** Associations between age participation coefficients across cortex. **(D)** Associations between play and participation coefficients across cortex. For panels A-D, the upper rows show results for all brain parcels, and the lower rows display uncorrected results with *p* < 0.05.

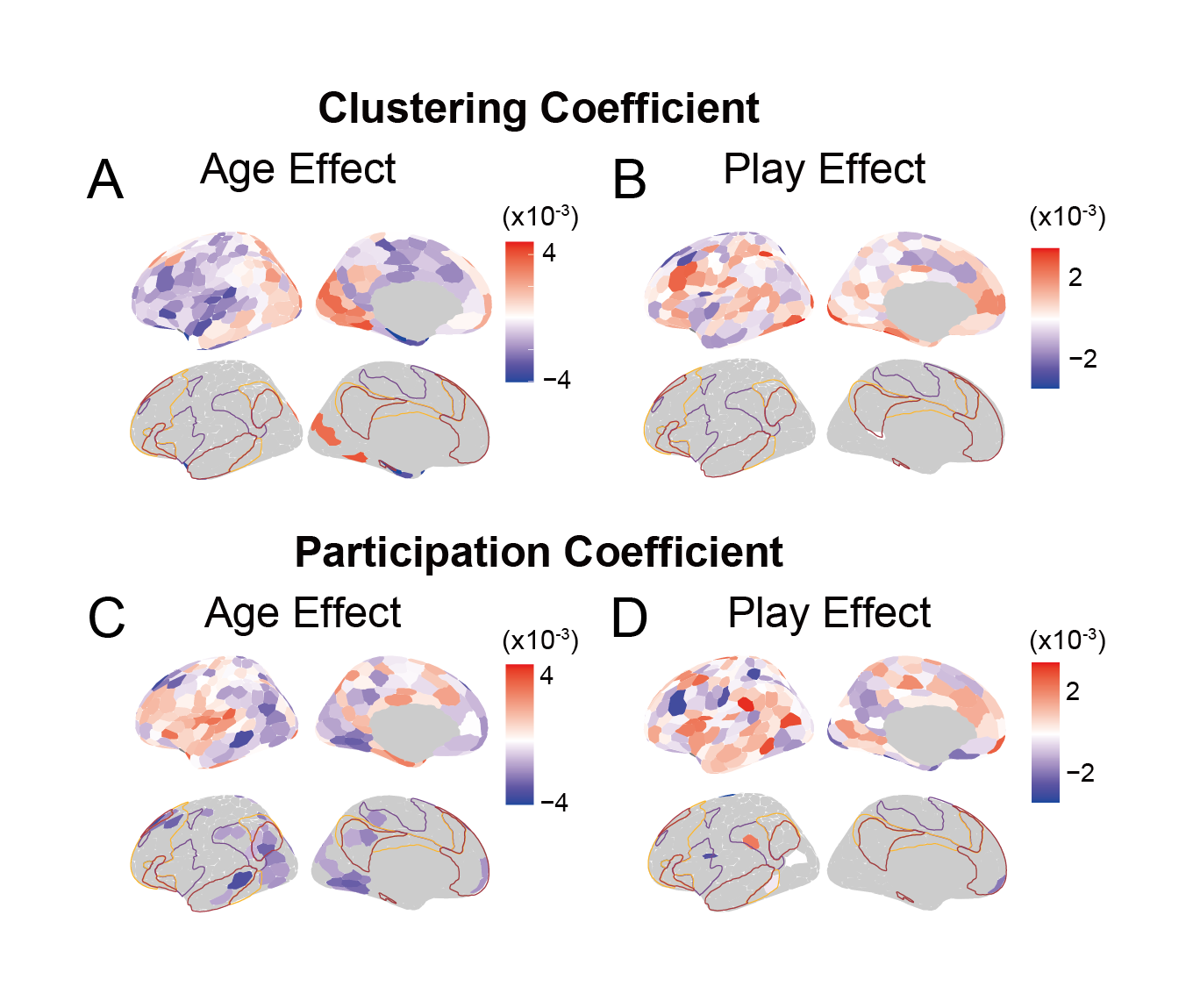

**Supplementary Figure 5**

*Associations between age, play and functional network measures in the left hemisphere in childhood*

### Supplementary Materials: Study 3

**Supplementary Table 3**

*Associations with age in the Montessori dataset*

|  | Montessori group | | | Traditional group | | |
| --- | --- | --- | --- | --- | --- | --- |
|  | *R^2^* _Partial_ | *P* | *P_FDR_* | *R^2^* _Partial_ | *P* | *P_FDR_* |
| DMN-DMN | 0.179 | 0.144 | 0.697 | 0.001 | 0.983 | 0.983 |
| DMN-CN | 0.016 | 0.584 | 0.938 | 0.023 | 0.508 | 0.612 |
| DMN-SAL | 0.001 | 0.988 | 0.988 | 0.029 | 0.448 | 0.612 |
| CN-CN | 0.073 | 0.232 | 0.670 | 0.079 | 0.286 | 0.612 |
| CN-SAL | 0.011 | 0.650 | 0.938 | 0.022 | 0.510 | 0.612 |
| SAL-SAL | 0.004 | 0.782 | 0.938 | 0.050 | 0.317 | 0.612 |

**(A–F)** GAM-predicted developmental trajectories accompanied by a 95% confidence interval of DMN–DMN (A), DMN–CN (B), DMN–SAL (C), CN–CN (D), CN–SAL (E), and SAL–SAL (F) functional connectivity in children attending traditional schools (gray) and Montessori schools (purple).

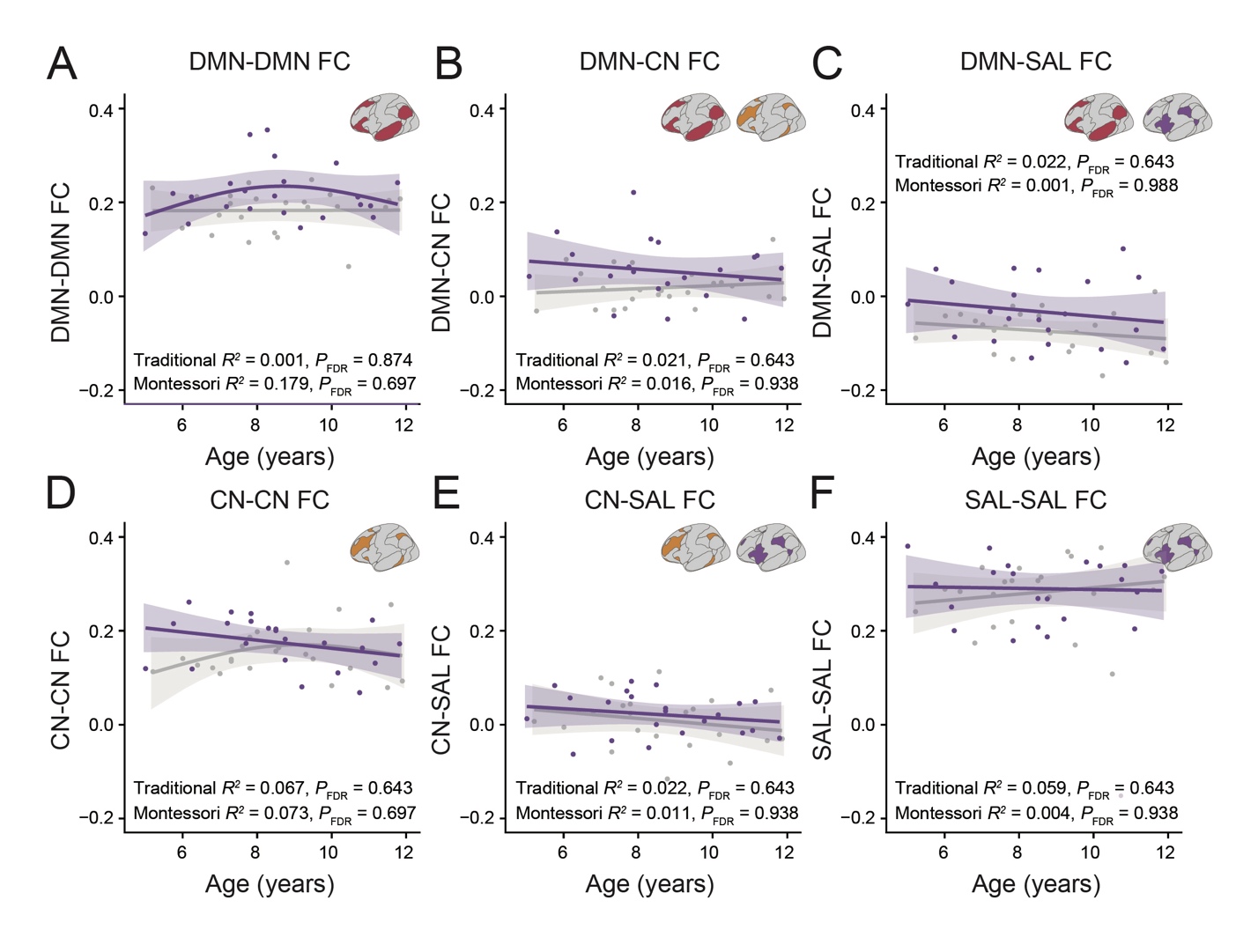

**Supplementary Figure 6**

***Development of brain functional connectivity by school environment***

**Supplementary Figure 7**

*Sensitivity analysis*

**(A)** SES composite score per by school environment (*t* = 1.742, *P* = 0.089). **(B)** Heatmap map of correlational strength between functional connectivity and SES composite score. **(C)** Heatmap map of correlational strength between functional connectivity and school environment controlling for SES. * for uncorrected *p* < 0.05.

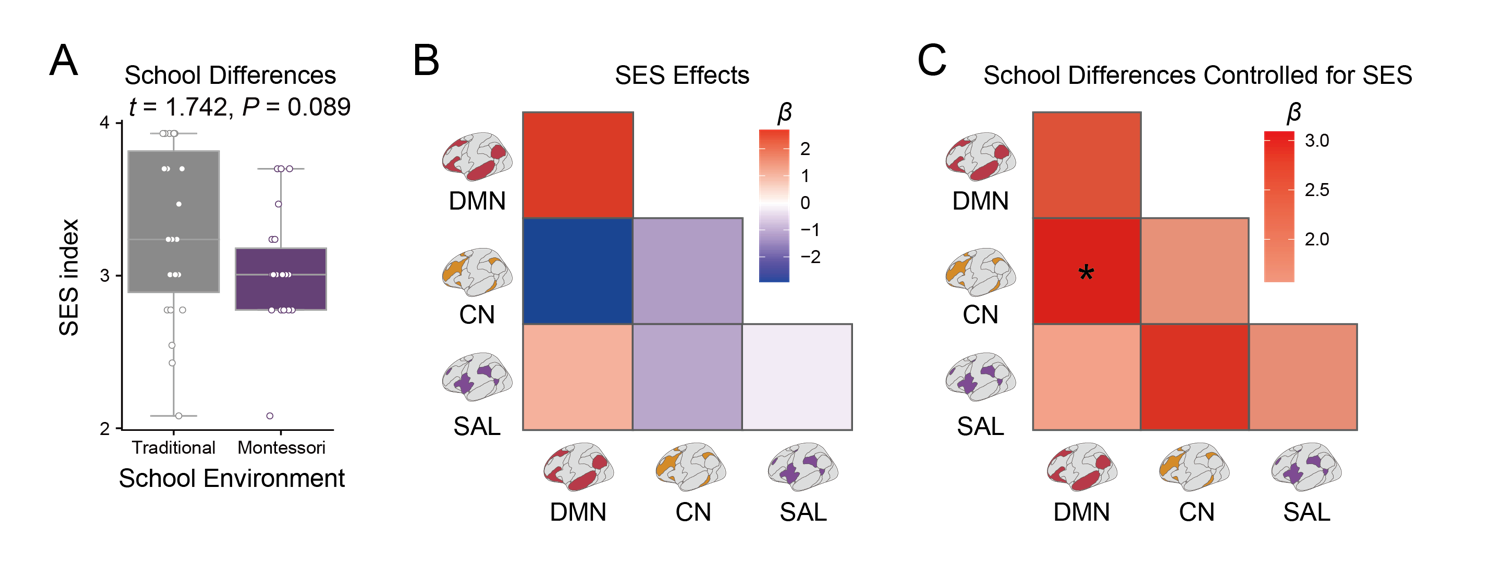
